# Context-dependent signaling of coincident auditory and visual events in primary visual cortex

**DOI:** 10.1101/258970

**Authors:** Thomas Deneux, Alexandre Kempf, Brice Bathellier

## Abstract

Detecting rapid coincident changes across sensory modalities is essential to recognize sudden threats and events. Using two-photon calcium imaging in identified cell types in awake mice, we show that auditory cortex (AC) neurons projecting to primary visual cortex (V1) preferentially encode the abrupt onsets of sounds. In V1, a sub-population of layer 1 interneurons gates this selective cross-modal information by a suppression specific to the absence of visual inputs. However, when large auditory onsets coincide with visual stimuli, visual responses are strongly boosted in V1. Thus, a dynamic asymmetric circuit across AC and V1 specifically identifies visual events starting simultaneously to sudden sounds, potentially catalyzing localization of new sound sources in the visual field.

## Introduction

Numerous multisensory illusions^1–4^ show that audition and vision have strong perceptual bonds. For example, in the double flash illusion^3^ a brief sequence of two sounds played simultaneously to a single visual flash leads to the impression of two flashes. While the mechanisms of cross-modal perceptual interactions remain unclear, the anatomy of the mammalian brain shows multiple sites of auditory-visual convergence. One of the best studied examples happens in the superior colliculus, in which visual and auditory cues are combined in the networks computing gaze direction^5,6^. In cortex, secondary associative areas, such as parietal cortex^7,8^, are not the sole cortical sites of auditory-visual convergence: increasing evidence shows that functional auditory-visual interactions exist already in primary sensory cortex^9–13^. Moreover, axonal tracing studies indicate that significant direct connections exist between primary auditory and visual cortex^14–20^, suggesting that important auditory-visual computations are implemented before the associative stage. The role of these direct connections has started to be addressed in the mouse model. Recent studies indicate that, in mice, auditory to visual connections are much stronger than their reciprocals, and that they provide very significant inputs to visual cortex that can modulate visually-driven activity^11,12^. Nevertheless, the computational role of early auditory-visual connections remains unclear, in particular because information is lacking about the auditory features channeled from auditory to visual cortex and how they are combined with the visual processing stream.

Mouse auditory cortex encodes a wide variety of acoustic features^21^ ranging from sound frequency^22,23^ to temporal features such as modulations of frequency^24^ or intensity^25^ including the salient intensity variations occurring at sound onsets and offsets^26,27^. The question of how sound frequency information would map onto visual cortex is a difficult one, because of the lack of perceptual and ethological data on the particular frequency cues that could potentially be associated with particular visual stimuli. In contrast, temporal information is known to be used for perceptually assigning auditory and visual stimuli to the same object, based on temporal coincidence as in the double flash^28^ and ventriloquist illusions^1,29^ and on covariations of particular features such as the size of the visual input and of the sound intensity envelope as classically evidenced with looming and receding auditory-visual stimuli^30^. Yet it is not known what time and intensity features are channeled through the direct connection between AC and V1 and how they impact visual processing.

To address this question, we here used two-photon calcium imaging and intersectional genetics in order to identify the particular intensity envelope features encoded by AC neurons that project to V1. We show that V1-projecting neurons preferentially encode a subset of the intensity variations extracted by auditory cortex, with a strong enrichment in cells sensitive to the onset of steeply rising sounds. Strikingly, we also show that this selective information impacts V1 in an illumination-dependent manner, with a net inhibitory effect in darkness and a net positive effect in the light, thus reconciling opposing observations reported previously^11,12^. Analyzing the responses of V1 excitatory and inhibitory neurons in layer 1 and layer 2/3, we show evidence that inhibition in darkness originates from a specific subpopulation of L1 inhibitory cells which masks the direct excitatory drive provided by AC axons to pyramidal cells^12^. Interestingly, the activity of this L1 subpopulation is reduced in normal light condition releasing the excitatory drive from AC. Furthermore, we show that this mechanism also allows for a specific boosting of visual responses which occurs together with large and rapid auditory onsets, and that this boosting correlates with improved detection of visual stimuli in a behavioral task. We thus propose that one important role of auditory to visual cortex connections is to increase saliency of visual events that occur simultaneously to the rise of a sudden sound source, possibly to help identifying the particular visual events responsible for new sounds.

## Results

### V1-projecting neurons in auditory cortex preferentially encode salient sound onsets

Intensity variations are precisely coded in auditory cortex, in particular the direction of variations which is clearly reflected in so called “On” (increasing intensity) or “Off” (decreasing intensity) responses^26,27,31,32^, but also in neurons that more tonically follow sound amplitude^27,33^. We have recently shown that not only the direction but also the amplitude of variations is coded in different cell types. For example, “Loud On” neurons specifically respond to high amplitude sound onsets, while “Quiet On” only respond to low amplitude (and not high amplitude) onsets^27^. The advantage of this coding scheme is to better tile the possible range of variations and thereby refine sound categories that reflect different types of events. For example, sounds coming from sudden physical events such as shocks tend to rise abruptly in intensity and rather activate “Loud On” neurons. On the contrary, sounds coming from more continuous events slowly ramp up in intensity and thus first activate “Quiet On” neurons. Ideal stimuli to capture the variety of On, Off and more tonic response types are long up- and down-ramping intensity profiles. We thus wondered whether such stimuli are similarly encoded by generic auditory cortex neurons and by neurons projecting to the visual cortex. To identify the latter, we injected in V1 a canine adenovirus (CAV) expressing Cre, which is retrogradely transported through axons. In the same mice, we injected in AC an adeno-associated virus (AAV) expressing GCAMP6s^34^ in a Cre-dependent manner (**Fig. 1a**). As a result, GCAMP6 expression was obtained exclusively in AC neurons that project to V1. Consistent with previous reports we observed that these neurons were located predominantly in layer 5 (**Fig. 1b**)^12^. As a comparison, in another set of mice, we injected an AAVl-syn-GCAMP6s virus to broadly express the calcium reporter in generic auditory cortex neurons. Using two-photon microscopy, we imaged calcium responses at single cell resolution in these two set of mice (layer 2/3: 3771 neurons, 18 sessions, 7 mice; V1-projecting: 1593 neurons, 14 sessions, 3 mice) while playing white noise sounds ramping up or down in intensity (range 60 to 85dB – note that visual stimuli were also played in these experiments but had no impact on AC activity, **Supplementary Fig. 1**).

**Figure 1:**
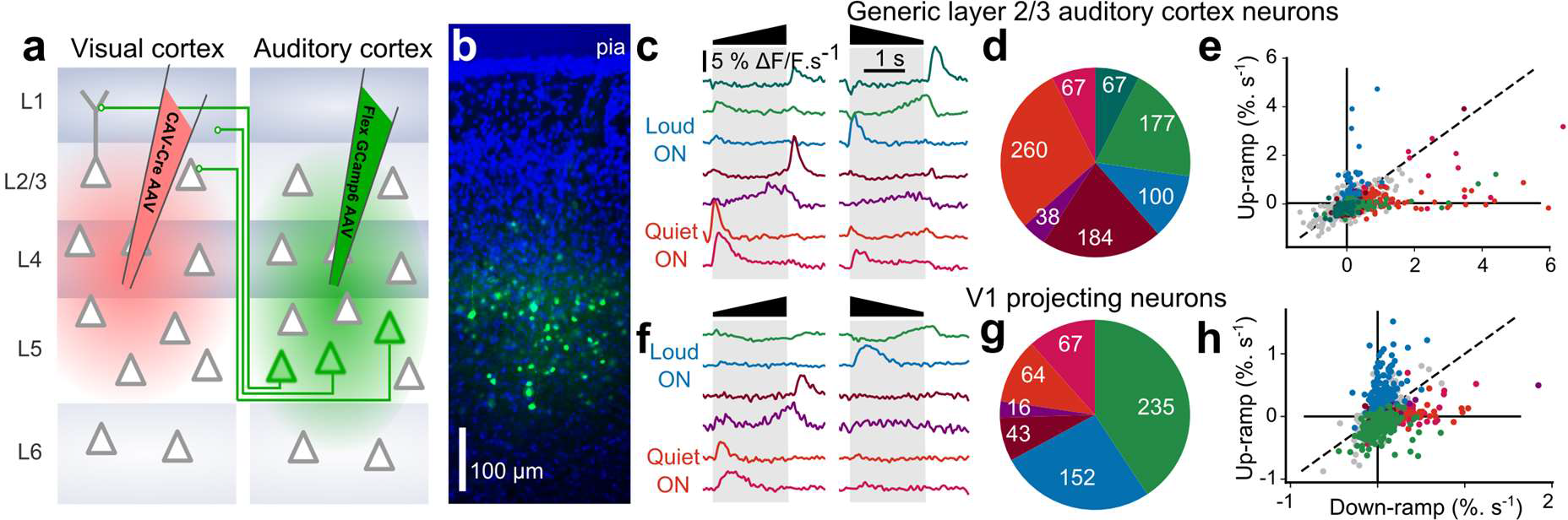
V1-projecting neurons over-represent loud onsets of sounds. **a.** Viral expression strategy for GCAMP6s labelling of V1 projecting neurons in AC. **b.**Epifluorescence image of an AC histological section (blue = DAPI, green = GCAMP6s) showing V1 projecting neurons (cortical layers are matched to **a**). **c.** Response profiles and cell count distribution of the 7 different activity clusters fund in layer 2/3 AC neurons. Color code: Blue and greens = clusters preferring down ramps, warm colors = clusters preferring up-ramps. **d.** Fraction of neurons corresponding to each cluster shown in c, with the absolute number neurons superimposed. **e.** Distribution of the mean response to up-and down-ramps for the neurons included in the responses clusters shown in c. **f-h.** Same as **c-e** but for the 6 clusters identified in V1 projecting neurons.

After automated ROIs extraction^35^, calcium signals were deconvolved^36,37^ and smoothed to obtain a temporally more accurate estimate of actual neuronal firing rates. The resulting averaged response profiles were then submitted to a hierarchical clustering^27^ (see Methods) to identify the different types of responses. In generic AC neurons, we found 7 distinct clusters, unsurprisingly corresponding to “On”, “Off” or tonic response types, each with a preference for either lower or higher sound amplitudes (**Fig. 1c**). Also, as reported previously^27^, a majority of neurons preferred up-ramps (62% of clustered neurons, **Fig. 1c**). Interestingly, in V1 projecting neurons we identified the same response clusters, to the exception of the sparser “Quiet Off” responses (**Fig. 1d**), however, the distribution of each response types was clearly different. First, in contrast to generic L2/3 neurons, a majority of V1-projecting neurons preferred down ramps (67% of clustered neurons, p<0.0001, bootstrap, see Methods **Fig. 1d**). Second, some cell types that were rare in the generic L2/3 AC population were clearly enriched in V1 projecting neurons. Most prominently, a 2.5-fold increase was observed for "Loud On" responses (10% of generic clustered cells, and 26% of clustered V1 projecting neurons, p<0.0001, bootstrap, **Fig. 1c,d**). Thus, while V1-projecting neurons encode a broad range of intensity variation features, they clearly emphasize a subset of them, in particular sudden onsets.

### The sign of auditory responses in V1 depends on the illumination-context

How do these specific AC inputs impact V1? One study in which mice received no visual input (dark environment) suggests an inhibitory effect^11^ while another study in which mice received visual stimuli describes excitatory effects^12^. Thus, the sign of AC inputs to V1 pyramidal cells could be illumination-dependent. To address this question, we performed two-photon calcium imaging in head-fixed awake mice (**Fig. 2a)** using the calcium sensor GCAMP6s (**Fig. 2b**) expressed in V1 through stereotaxic injection of an AAV-syn-GCAMP6s viral vector. Imaging of the same neurons was performed either in complete darkness or with a visual context, in the light, in front of a grey screen at low luminance (0.57 cd/m^2^; note that when screen is on, the mouse also receives visual inputs from the surrounding of the screen). To monitor gaze stability, pupil position and diameter was tracked during the experiment (**Fig. 2c,d, Supplementary Fig. 2**). V1 was identified using Fourier intrinsic imaging^38^ as the largest retinotopic field in visual areas (**Fig. 2e,f**), and all two-photon imaging fields-of-view were mapped to the retinotopic field thanks to blood vessel landmarks (**Fig. 2e,f**). All neurons imaged outside V1 according to this criterion (**Fig. 2f)** were analyzed separately. As visual sampling can be influenced with auditory stimuli^5^, we first measured whether sounds impacted triggered eye-movements. We found that sounds and in particular the sharp onsets of down-ramps triggered occasional and mostly horizontal eye saccades (**Fig. 3a**). A sound induced change in pupil diameter was also observed (**Fig. 3b, Supplementary Fig. 2**). In the light, the saccades triggered responses in V1 likely reflecting visual inputs as these responses vanished in the dark (**Supplementary Fig. 2**). Therefore, we excluded all trials with a saccade larger than mouse visual acuity (2° of visual angle) from our analyses (**Fig. 3a**). After this correction and pulling together the activity of 18925 V1 neurons (35 sessions, 9 mice), we observed that sounds alone trigger responses in supragranular V1 neurons, and that, as expected from the activity of V1-projecting AC neurons, these responses were stronger for the down-ramp (**Fig. 3b,c**).

**Figure 2:**
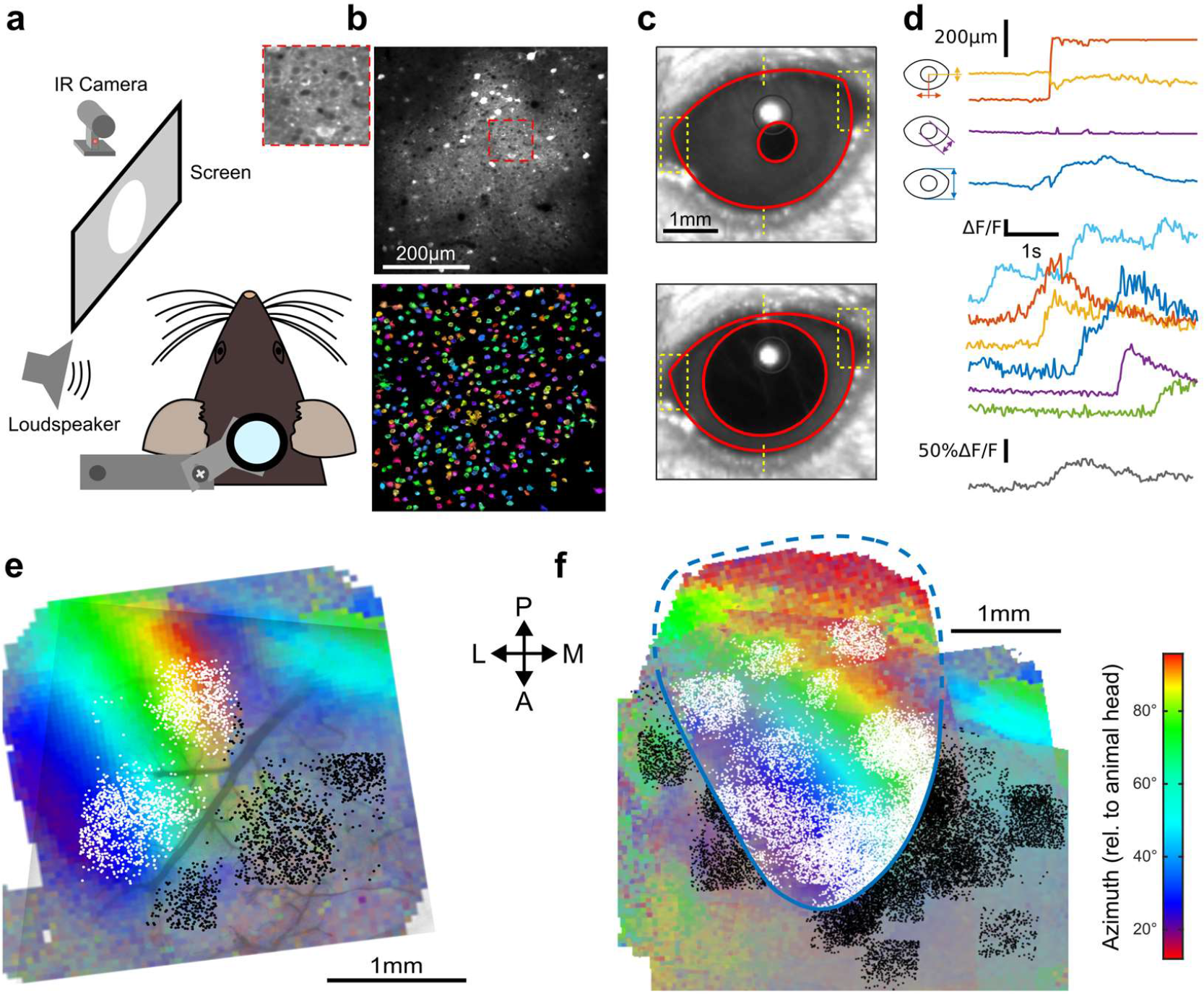
Retinotopically-mapped two-photon imaging fields in V1. **a.** Sketch of the experimental setup. **b** (top) Example of a 0.5×0.5 mm imaging field of view showing GCAMP6s-expressing neurons in V1 (top with magnification in inset) and the ROIs that were automatically detected as putative neurons (bottom, see Methods). **c.** Examples of eye tracking images. **d.** (top) Eye tracking showing a large saccade during a blank trial. The blue, purple and yellow-red traces indicate apertures between eye lids, pupil diameter, x-y motion ofpupil center is shown in red. (bottom) Examples of raw GCAMP6s traces from individual neurons recorded concomitantly in V1 (black, frame rate: 31Hz) and population average (grey) showing saccade-related neuronal activity. **e.** (left) Example of a retinotopic map obtained with Fourier intrinsic imaging (see Methods) and (right) registered maps across 9 mice. The color code indicates the azimuth in the visual field.

**Figure 3:**
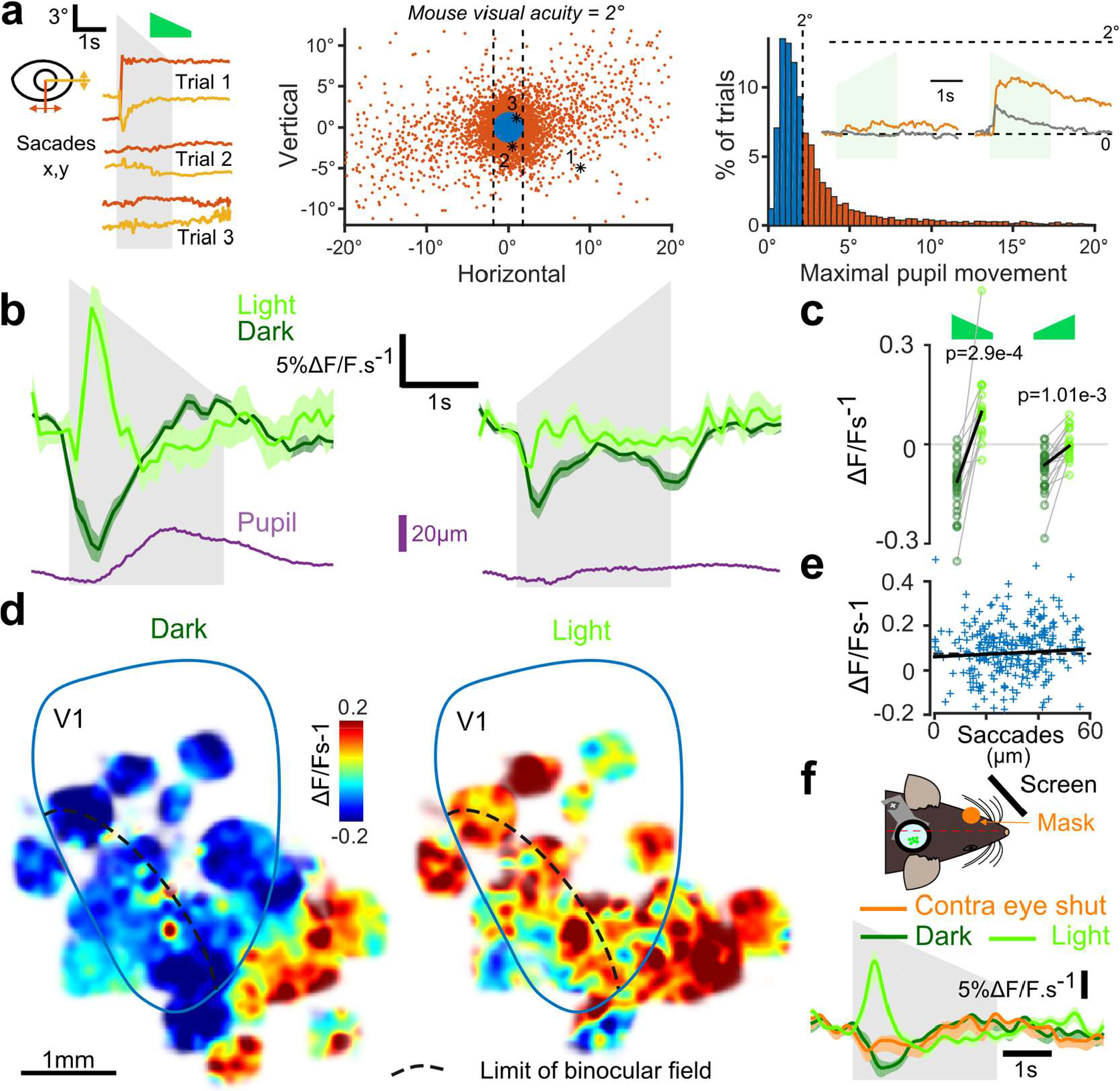
Auditory responses in V1 switch sign with the presence of visual inputs. **a.** (left) Horizontal and vertical pupil movements recorded during three trials. (center and right) Distribution of saccades across all mice (n = 7) and trials. The blue portion indicates trials where eye pupil movement did not exceed visual acuity in the mouse (2° of visual angle: 84% of all trials). Inset: average saccade responses to up-ramps and down-ramps, before (orange) and after (grey) trial filtering. **b.** Averaged deconvolved calcium traces of V1 neurons in the light (light green) and in the dark (dark green) (6207 neurons, n = 17 sessions in 7 mice). The purple line is the average pupil diameter. **c.** Average V1 response are larger in the light than in the dark (n = 17 recording sessions in 7 mice, Wilcoxon signed rank test). **d.** Smoothed maps of the local responses to the down-ramping sound, averaged across sessions and animals after registration with respect to the retinotopic map. **e.** Single trial saccade amplitudes (below the 2° visual acuity threshold) do not correlate with the amplitude of V1 population responses to sounds (Pearson correlation coefficient 0.05, p = 0.42). **f.** Mean V1 responses (2226 neurons, n = 13 sessions in 2 additional mice) in the dark, in the light and with the contralateral eye reversibly occluded as depicted in the sketch above.

In addition, we clearly observed that the net population response was globally inhibitory when mice were in complete darkness and excitatory when mice were in the light (**Fig. 3b,c**), corroborating the idea that the sign of AC inputs depends on the visual context (darkness vs visual scene). To test whether this context-dependency was a general property of AC outputs or if it was specific to AC inputs impacting V1, we measured auditory responses in the secondary visual or associative areas next to V1 which also receive inputs from AC^19^. Interestingly, these areas displayed excitatory responses both in the light and in the dark (**Fig. 3d**). Moreover, they showed richer auditory information, including selectivity to frequency (in FM sweeps) and to sound offsets, suggesting that they perform different cross-modal computations than V1 (**Supplementary Fig. 3**). As a control, we verified that all excitatory effects in the light were not due to unfiltered micro-saccades as the amplitude of these extremely small saccades was not correlated to sound responses (**Fig. 3e**). Moreover, excitatory responses to sounds could not be attributed to sound-induced pupil dilation as this effect occurred after V1 responses (**Fig. 3b**). We then wondered whether the context dependence was related to a global change in arousal caused by complete darkness which would involve both hemispheres simultaneously, or to the absence of inputs to the visual system, which could be induced unilaterally. We therefore repeated the experiments with masking of the contra-lateral eye and observed that this condition reproduced the responses of the dark context (**Fig. 3f**), ruling out the possibility of a global arousal effect. Thus, altogether we concluded that V1 implements a specific mechanism that reverses the impact of AC inputs depending on whether or not visual inputs are available.

### Minimal model for the context-dependence of auditory responses

To better apprehend the circuit mechanisms of this context-dependence, we looked for a minimal model of the effect, based on the facts that AC projections excite both pyramidal cells and interneurons in V112, and that V1-projecting AC neurons are not affected by light (Supplementary Fig. 1).

Considering a generic population of L2/3 V1 neurons (E-neurons), the minimal circuit which can provide sound-dependent inhibition to this population is a connection from an inhibitory population (I-neurons) driven by AC inputs (**Fig. 4a**). In this case, as we exemplified in a simple simulation (**Fig. 4b,c**), to render this inhibition context-dependent, the I-neurons must have (i) a non-linear response curve (threshold function), and (ii) must be inhibited in the presence of visual inputs (light modulation, **Fig. 4a**). Then, if I-neurons are close to activation threshold in the dark, and if inhibition by visual inputs brings them well below threshold in the light, I-neurons will be less active in the light than in the dark, leading for E-neurons to a dominance of excitation in the light and of inhibition in the dark (**Fig. 4b,c**). This simplistic model could be extended into more complex models in which inhibition of I-neurons in the light is more explicitly implemented, but as we show in **Supplementary Fig. 4**, whatever the complexity of the model, the context-dependent sign of auditory response in V1 requires the existence of neurons that have the typical response signature of I-neurons in light and dark **(Fig. 4d)**.

**Figure 4:**
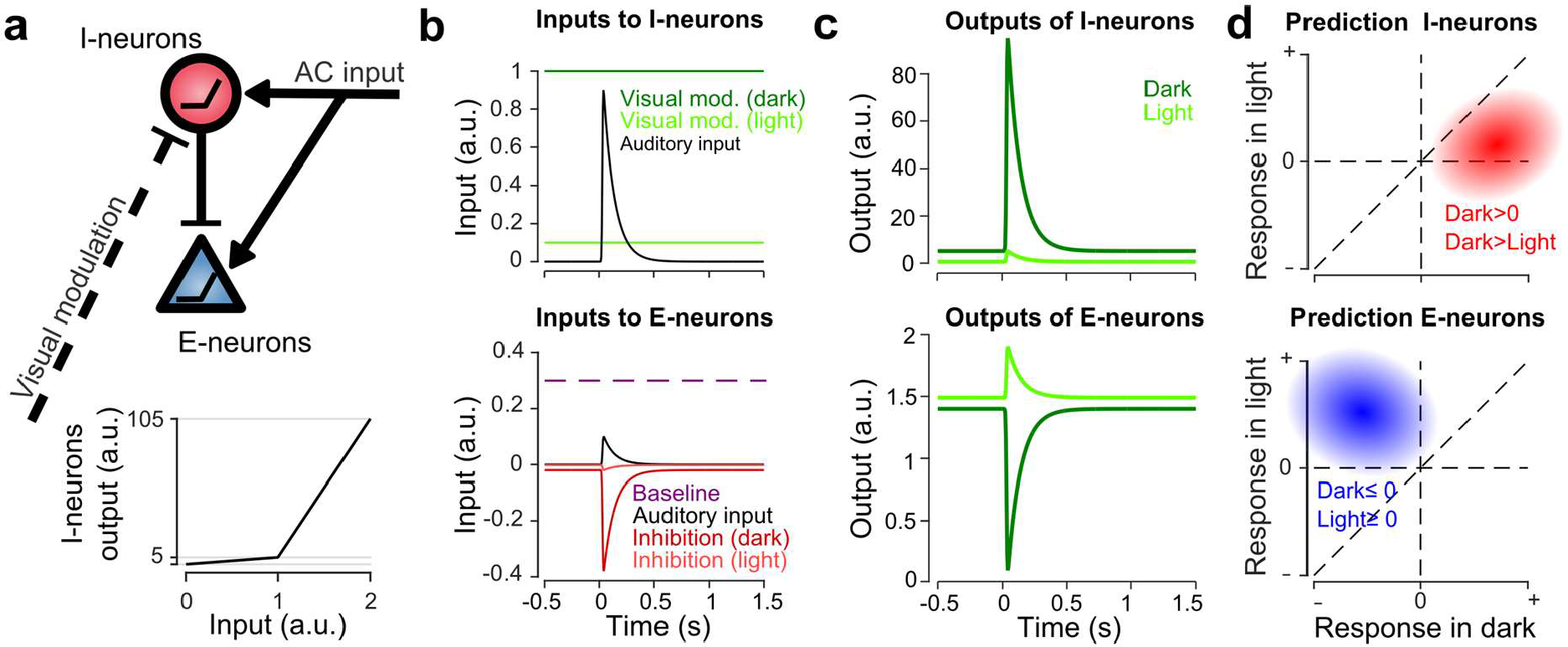
A minimal model predicts the response signature of neurons mediating sound-induced inhibition. **a.** Sketch of the minimal model for the switch between negative and positive sound responses in dark vs in light. An inhibitory population (I-neurons) endowed with a non-linear response function receives positive auditory and negative visual inputs. If both inputs are present the drive is below threshold and no inhibition is delivered. Thus, only direct excitatory inputs impact the main neuronal population (E-neurons). **b.** Input currents delivered to the I- and E-neurons during the simulations. **c.** Outputs of the two populations in the presence (light) and absence (dark) of visual input. **d.** Predicted responses signatures for E- and I-neurons as defined in the model.

### Layer 1 contains an interneuron subpopulation delivering context-dependent inhibition

We thus aimed at identifying the predicted interneuron population that mediates context-dependent inhibition in V1. Because anatomy shows that AC to V1 projections are concentrated in L1 and L2/3^12^, we imaged neurons in these two layers broadly labeled with GCAMP6s (synapsin promotor) in GAD2-Cre x flex-TdTomato mice which allow identifying inhibitory neurons based on red fluorescent protein expression (**Fig. 5a**). We observed that in layer 2/3 both excitatory (n=4348) and inhibitory neurons (n=726) were globally inhibited by sounds in the dark and excited in the light, while the layer 1 interneuron population seemed globally unaffected in the dark and excited in the light (**Fig. 5b**). However, the population trend concealed functionally distinct subpopulations. When we tested for significant positive or negative responses in the dark in single neurons (Wilcoxon rank-sum test, p<0.01), we found a large fraction of neurons significantly inhibited, but also, a subpopulation of 12.8% of all L1 inhibitory neurons that were significantly activated in the dark (**Fig. 5c**). This positive response was not observed in L2/3 above significance threshold (**Fig. 5c**). When plotting the response of L1 neurons in the dark against in the light (**Fig. 5d**), it also became apparent that the L1 neurons that are excited by sounds in the dark tend to be less excited in the light (e.g. **Fig. 5e**). In contrast, L1 neurons inhibited or unaffected by sounds in the dark became more activated in the light (**Fig. 5d**). Consistently, statistical assessment showed that a significant fraction of 5.3% of L1 neurons (Wilcoxon rank-sum test, p<0.01) respond less in light than in dark (**Fig. 5f**), and that almost all of these neurons are significantly excited in the dark (4.9%, **Fig. 5g**). In contrast, based on the same tests, there was no significant population of L2/3 neurons that responded less in light than in dark (**Fig. 5f, g**). Thus, we concluded that layer 1 contains the only supra-granular subpopulation of GABAergic neurons that can provide a sound-induced inhibition gated by visual inputs. These neurons are thus good candidates to mediate the context-dependence of sound responses observed in the bulk of L2/3 V1 neurons (**Fig. 5h**).

**Figure 5:**
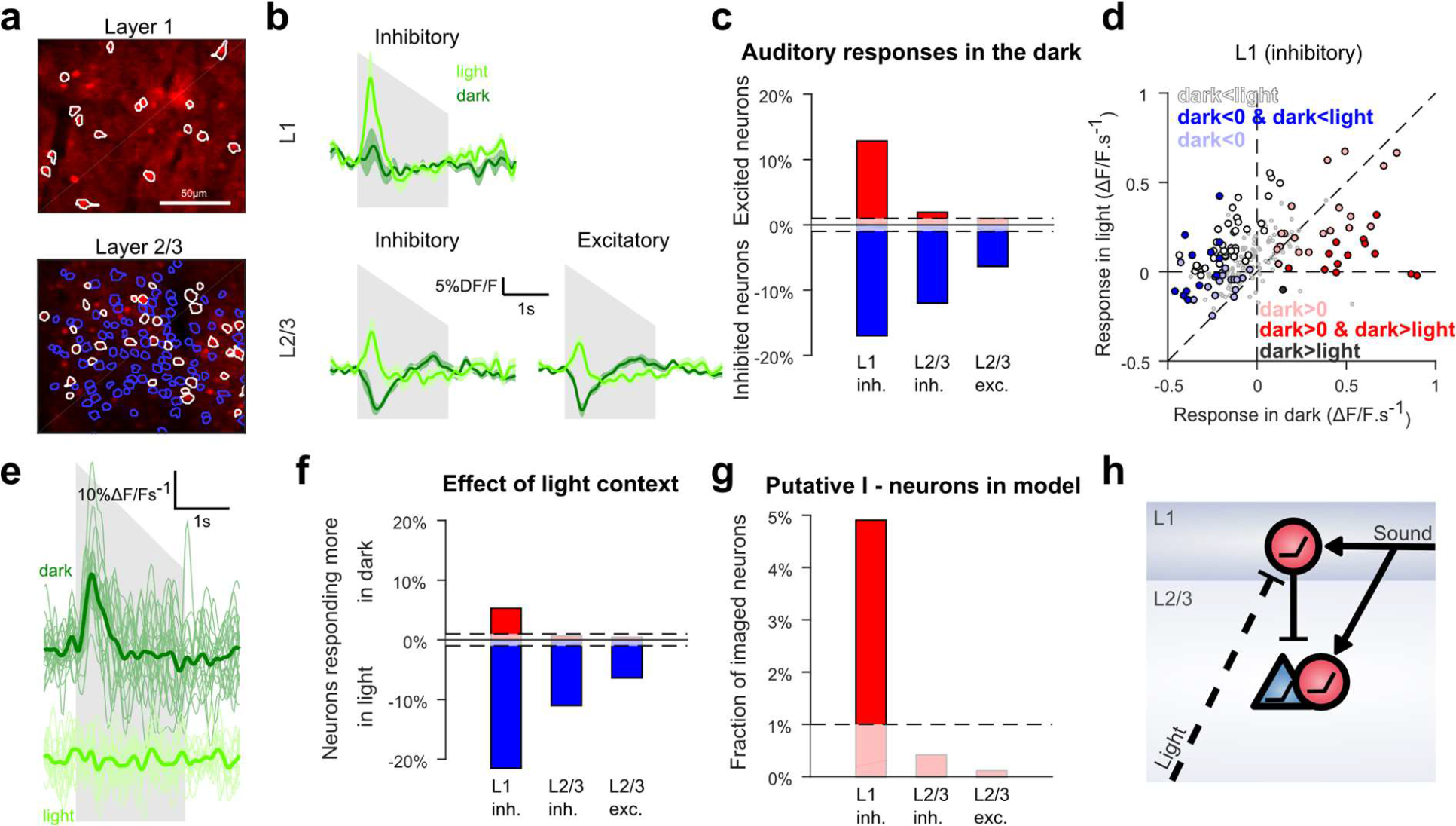
Context-dependent inhibition by sound is mediated by a L1 subpopulation. **a.** In vivo 2-photon images of GAD2-positive V1 neurons in L1 and L2/3 expressing td-Tomato. Superimposed are the contours of the active regions of interest identified by GCAMP6s imaging. **b.** Mean responses to the down-ramping soundfor inhibitory and excitatory neurons in L2/3 and L1. **c.** Fraction of neurons significantly excited (red) or inhibited (blue) by sounds for each layer and cell type (Wilcoxon rank sum test, p<0.01). **d.** Scatter plot of the mean responses of all L1 inhibitory neurons to the down-ramping sound in the light against in the dark. Small grey dots indicates neurons that do not significantly respond in any condition. Larger, colored dots indicate significantly responding neurons. Dark blue indicates neurons significantly inhibited in the dark and responding less in dark than in light. Dark red dots indicates neurons significantly activated in the dark and responding more in dark than in light. Neurons responding equally in dark and light are marked with light blue (negative response) and light red (positive response) dots. Neurons that are non-responsive in the dark but respond more or less in the light are mark in white and dark respectively. **e.** Single trial deconvolved traces for an L1 interneuron active in the dark and inactive in the light. **f.** Fractions of neurons cell type responding significantly more in light than in dark (blue) or in more in dark than in light (red) (Wilcoxon rank sum test, p<0.01). **g.** Fraction of neurons that can mediate context dependent inhibition in each layer i.e. interneurons significantly excited by sounds in the dark and responding significantly less in the light than in the dark. **h.** Schematics of the core mechanism for context-dependent auditory responses in V1.

### Sounds with large onsets boost the representation and perception of coincident visual events

We then assessed the impact of AC inputs to V1 on the representations of visual events. We recorded 9849 L2/3 neurons (23 sessions, 7 mice) expressing GCAMP6s under the synapsin promotor while delivering 2 sec long up- or down-ramping auditory stimuli together with a white disk looming or receding coincidently over the same duration as the sounds. The loudspeaker was placed such that auditory and visual stimuli came from a similar direction. Unimodal stimuli were also delivered to assess additivity for the bimodal conditions. We first observed that many V1 neurons displayed a supra-additive boosting of their visual response when coincident with the large onset of down-ramps (e.g. **Fig. 6a**), an effect even visible at the population level (**Fig. 6b**). Because looming and receding stimuli trigger different type of visual responses in V1, we performed a clustering in order to identify, in a model-free manner, the main types of visual responses and sound-induced modulations (see Methods, note that only the V1 neurons with high signal-to-noise and sufficient number of trials without eye movement, n=499, were retained by the clustering). This analysis identified seven distinct clusters (**Fig. 6c**), among which three clusters (254 neurons) responded more to looming than receding disks, and three clusters (141 neurons) had the opposite preference. Four clusters were tightly direction-specific and responded only for their preferred stimulus, at its beginning or end, while the three others were less specific to direction, but also displayed temporal specificity. The temporal specificity was probably due to the location of the retinotopic receptive field of the neurons with respect to stimulus centre.

**Figure 6:**
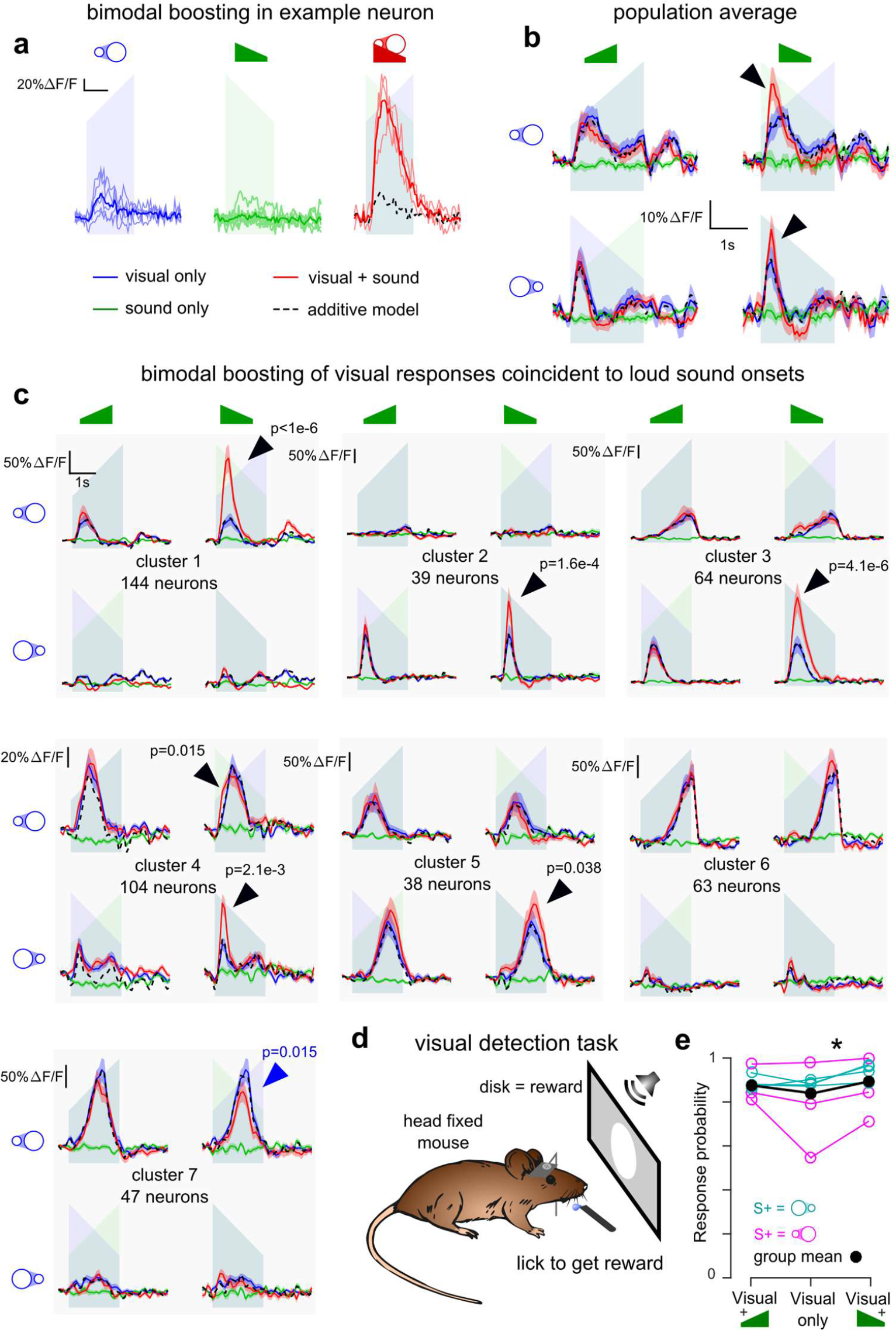
V1 neurons are boosted by the coincidence of large auditory onsets with visual onsets. **a** Examples of multisensory neurons in V1. Raw GCAMP6s traces for a visual stimulus (blue), a sound (green) and a combination of both (red) show a large neuronal response for bimodal condition compared to additive prediction (dashed black line) based on unimodal responses. Individual trials (thin lines) and their average (thick lines) show that the multisensory responses are robust for those neurons. **b.** Average deconvolved calcium responses (mean ± sem) in V1 for up- and down-ramping sounds (green), looming and receding visual stimuli (blue) and their four bimodal combinations. Black dashed lines: linear prediction. **c.** Uni- and bimodal responses, displayed as in b., of the six different functional cell types identified by hierarchical clustering. **d.** Sketch of the behavioral experiments. **e.** Increase of visual stimulus detection rate when a coincident down-ramping sound is played (Wilcoxon signed rank test, p = 0.0156); a non-significant increase is also observed with the up-ramping sound (p = 0.375).

The direction specificity likely reflects the direction specificity of their receptive field, although we do not exclude that some of the specificity also relates to the global geometry of the stimulus as direction selectivity measured in the same neurons with drifting gratings was much less sharp than with receding/looming disks (**Supplementary Fig. 5**). Also many cells responded to the disk but not to the gratings and conversely, suggesting complex nonlinearities in receptive fields (**Supplementary Fig. 5**). But more interestingly, the bimodal conditions revealed very clear supra-additive responses. Strikingly, all clusters responding at the onset of visual stimuli had their response strongly boosted by coincidence with the loud onset of down-ramps (**Fig. 6c,** black arrows for clusters 1-5, bootstrap test, see Methods). No or much weaker boosting was observed with the quiet onset of up-ramps (**Fig. 6c**), consistent with its reduced representation in AC to V1 projections (**Fig. 1**). Responses occurring towards the end of the stimulus were minimally (although significantly, cluster 5) boosted, probably because they correspond to auditory features (“loud and quiet tonic”) that are more weakly impacting V1 (see **Fig. 3b**). Our data thus suggests that visual responses in V1 are boosted specifically by the coincidence of loud sound onsets. Note that a moderate response suppression was also observed, but only in cluster 7 (**Fig. 6c**, blue arrow). Interestingly, the boosting of visual responses appeared to be a strong feature of V1, as the same analysis identified only one nonlinear cluster out of six clusters in the associative area medial to V1 (**Supplementary Fig. 6**).

We thus wondered whether V1 boosting could have a perceptual impact in mice. To do so, we trained head-fixed water-deprived mice to respond to looming or receding visual stimuli by liking at a water spout to get a reward (**Fig. 6d**). After training, mice reliably responded to the visual stimulus with a success rate of 83.7 ± 5.7 %. However, in trials in which the visual stimulus was paired with a down-ramp (loud onset) the response probability was significantly increased to 90.2 ± 4.1 % (**Fig. 6e**, Wilcoxon signed rank test, p = 0.0156). A weaker, nonsignificant effect was seen with up-ramps (**Fig. 6e**, 88.4 ± 2.2 %, Wilcoxon signed rank test, p = 0.375). Thus, loud sound onsets not only boost concomitant visual responses in V1, but also improve visual detection in a behavioral task.

### Audio-visual boosting can result from direct AC excitation if pyramidal cells are nonlinear

What could be the mechanism of sound-induced boosting? To answer this question, we attempted to extend our minimal model for sound responses in V1 (**Fig. 4**), in order to reproduce also the boosting effect. We realized, that the same model can reproduce boosting by sounds (**Fig. 7a–c**) provided that pyramidal cells have a simple non-linearity. We modeled this non-linearity with a thresholded activation function having a weak subthreshold output gain and a high supra-threshold gain (**Fig. 7a**). This non-linearity is necessary to account for the fact that responses to sound alone are much weaker than the bimodal boosting effects. With this design, we could also reproduce the observation that boosting mainly occurs for the preferred visual stimulus (**Fig. 6c**), provided that the auditory and non-preferred visual inputs are driving excitatory neurons in the subthreshold regime (**Fig. 7b**) while the preferred input brings excitatory neurons close or above threshold (**Fig. 7c**). In this case the auditory response sums with a high gain for the preferred stimulus and a low gain for the non-preferred. Thus altogether, our data and modeling work indicates that a minimal circuit including a direct excitation from auditory cortex onto L2/3 pyramidal cells and an indirect inhibition via a subpopulation of L1 interneuron is sufficient to provide an illumination-dependent auditory input that emphasizes visual events coincident to sound loud onsets.

**Figure 7:**
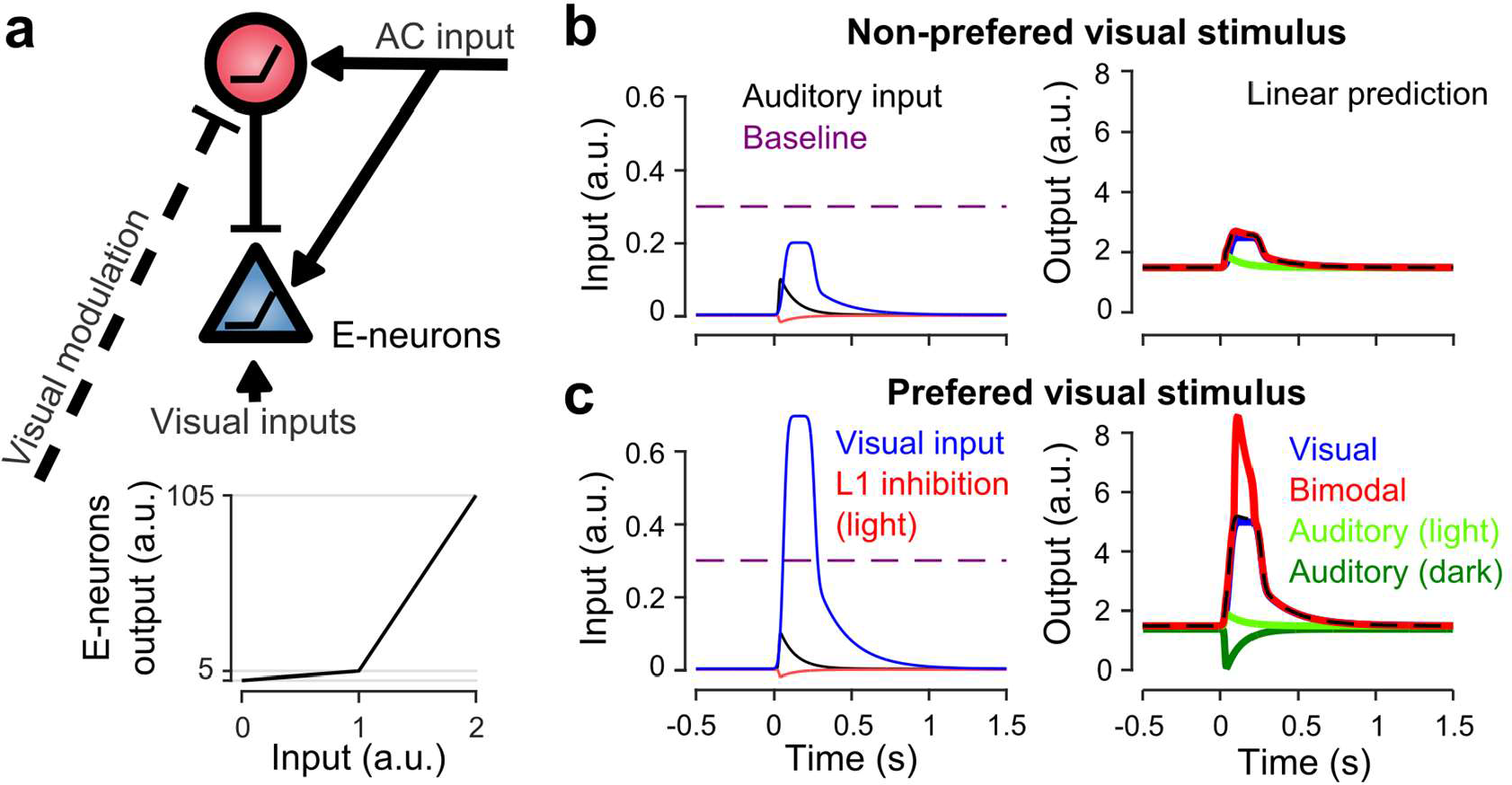
Sound-dependent boosting is reproduced by the L1 inhibition model. **a.** Sketch of gated inhibition model as in **Fig. 4a. b.** Input currents (left) received by the E-neurons population and output response (right) for a non-preferred, “subthreshold” visual input. **c.** Same as in b. but for a preferred, suprathreshold visual input.

## Discussion

Using two-photon calcium imaging in identified excitatory and inhibitory neurons during auditory and visual stimulation, we demonstrated three important features of AC to V1 connections. First, we showed that they preferentially transfer a subset of the temporal auditory features encoded by AC, most prominently large amplitude onsets. Second, they are gated by a context-dependent inhibitory mechanism likely implemented by a subpopulation of L1 interneurons. Third, their most salient impact on V1 representations is to produce a strong boosting of responses to dynamical visual stimuli concomitant to sharp sound onsets.

The preference of V1 projecting neurons as compared to bulk layer 2/3 neurons for the features of suddenly rising and slowly decaying sounds (**Fig. 1**) is an interesting novel case of coding bias in a projection-defined neuronal subpopulation of a cortical area. Similar results have been found in somatosensory cortex for neurons projecting to secondary sensory or motor cortex^39^. In AC, it is well established that specific acoustic features are related to the horizontal localization of neurons in different subfields^22,40,41^ and that, at least in higher mammals such as cats, these subfields can project to independent pathways propagating different acoustic information for behavioral decisions (e.g. localization and quality^42^). Our results show that projection specificity can also be a determinant of feature specificity, at least for intensity modulation features.

Strikingly, this select auditory information impacts V1 in a dynamic manner, with a net inhibitory drive in the absence of visual inputs and a net excitatory drive in layer 2/3 neurons in the presence of visual inputs (**Fig. 3b**). Sound-induced inhibition of V1 has been described earlier^11^. We here show that this inhibition is specific to darkness, and thus could specifically serve to decrease V1 activity in the dark when visual information is irrelevant for sound source localization. The illumination-dependence of auditory responses in V1 is reminiscent of the recent observation that AC to V1 projections, activated by loud sounds and acting through L1, boost preferred orientation responses in V1 and inhibit non-preferred orientations^12^. The two effects share several properties including selectivity to high sound levels, the dominance of inhibition for weaker visual inputs and of excitation for stronger visual inputs, and the involvement of L1 neurons for mediating inhibition. The main discrepancy is the apparent lack of visual input specificity of the illumination-dependent excitation of V1 neurons by AC inputs alone. This could be due to differences related to the type of visual inputs used. One important result appearing in our data is that only a fraction of all L1 interneurons have response properties compatible with the function of releasing an illumination-dependent, sound-triggered inhibition (**Fig. 5c,f,g**), while these properties are not found in L2/3 (**Fig. 5c,f,g**). Thus the silencing of visual processing by sounds in the dark is not a generic function of L1 interneurons, compatible with the observation that only VIP negative interneurons receive AC inputs in V1^12^ and the observation that L1 interneurons serve other function^43^, such as disinhibition of L2/3 in various contexts^44–48^. An interesting question that cannot be addressed with the current observations is how the specific L1 neurons that mediate sound-induced inhibition are themselves inhibited in the presence of visual inputs. Several putative mechanisms are compatible with the data including disinhibition (see models in **Supplementary Fig. 4**).

Another important phenomenon observed in our data is the strong boosting of visual responses coincident to loud sound onsets (**Fig. 6**). Our model suggests that this boosting results directly from the non-linear summation of visual and auditory inputs in V1 pyramidal cells, while sound-triggered inhibition is abolished by visual context (**Fig. 7**). The large amplitude of the sound-induced boosting as compared to the weaker responses observed when sounds are played without coincident visual input (**Fig. 3 and 5** *vs* **Fig. 6**, see also **Fig. 7**) points towards a threshold mechanisms, allowing for amplification of the auditory input only if sufficient visual input concomitantly arrives. Such a gated amplification mechanism could be implemented directly in L2/3 pyramidal neurons using the direct inputs they receive from auditory cortex^12,20^. For example, amplification of L1 input by coincident somatic inputs thanks to calcium spikes in the apical dendrite has been described both in L5^49^ and L2/3^50^ pyramidal cells. While this is a good candidate mechanism for the observed boosting effect, we cannot exclude alternative or complementary mechanisms such as a disinhibition mediated by a subpopulation of L1 neurons^12,46–48^, different from the subpopulation providing direct inhibition (**Fig. 5**). Also, as down ramping sounds are salient stimuli, a cholinergic input to visual cortex, known to increase cortical responses via desinhibition^44,45,51^ could partially contribute to the boosting effect. Independent on the mechanism, our data show that sound-induced boosting is a strong effect in V1 which tightly relates to coincidence with the onset of salient sounds. This effect provides a powerful way to highlight, in visual space, the visual events potentially responsible for the sound, and thus might be an essential element of the cortical computations related to the localization of sound sources^5^.

## Methods

### Animals

All animals used were 8-16 week-old male C57Bl6J and GAD2-Cre (Jax #010802) x RCL-TdT (Jax #007909) mice. All animal procedures were approved by the French Ethical Committee (authorization 00275.01).

### Two-photon calcium imaging in awake mice

Three to four weeks before imaging, mice were anaesthetized under ketamine medetomidine. A large craniotomy (5 mm diameter) was performed above the right primary visual cortex (V1) or the right auditory cortex. For the imaging of the auditory cortex, the right masseter was removed before the craniotomy. We then performed three to four injections of 150 nl (30 nl.min^-1^), AAV1.Syn.GCaMP6s.WPRE virus obtained from Vector Core (Philadelphia, PA, USA) and diluted x10. The craniotomy was sealed with a glass window and a metal post was implanted using cyanolite glue followed by dental cement. A few days before imaging, mice were trained to stand still, head-fixed under the microscope for 30–60 min per day. Then mice were imaged 1-2h per day. Imaging was performed using a two-photon microscope (Femtonics, Budapest, Hungary) equipped with an 8 kHz resonant scanner combined with a pulsed laser (MaiTai-DS, SpectraPhysics, Santa Clara, CA, USA) tuned at 920 nm. We used 20x or 10x Olympus objectives (XLUMPLFLN and XLPLN10XSVMP), obtaining a field of view of 500×500, or 1000×1000 microns, respectively. To prevent light from the stimulation screen to enter the microscope objective, we designed a ring silicon mask that covered the gap between the metal chamber and the objective. Images were acquired at 31.5 Hz during blocks of 5s interleaved with 3s intervals. A single stimulus was played in each block and stimulus order was randomized. All stimuli were repeated 20 times except drifting gratings (10 repetitions).

Drifting square gratings (2Hz, 0.025 cyc/°) of 8 different directions (0° from bottom to top, anti-clockwise, 45°, 90°, 135°, 180°, 225°, 270°, 315°), and a size-increasing (18° to 105° of visual angle) or decreasing white disk over a black background were presented on a screen placed 11 cm to the mouse left eye. The luminance of the black background and white disks was 0.57 and 200 cd/m^2^ respectively. The screen (10VG BeeTronics, 22 **x** 13 cm) was located 11cm from the contra-lateral eye, thus covering 90° **x** 61° of its visual field including some of the binocular segment. For auditory cortex recording, we used 250 ms constant white noise sounds at 4 different intensity modulations, and two up- and down-intensity ramping sound between 60 and 85 dB SPL. Note that microscope scanners emitted a constant background sound of about 45dB SPL. For visual cortex experiments, we used only the two intensity ramps and added two frequency modulated sounds, going linearly from 8 kHz to 16 kHz and vice versa. Up- and down-amplitude ramps were combined with the increasing and decreasing disks to form 4 multisensory stimuli. The loudspeaker was placed just next to the stimulation screen, facing the contralateral eye. All stimuli were 2 s long. Auditory and visual stimuli were driven by Elphy (G. Sadoc, UNIC, France). All sounds were delivered at 192 kHz with a NI-PCI-6221 card (National Instrument) an amplifier and high-frequency loudspeakers (SA1 and MF1-S, Tucker-Davis Technologies, Alachua, FL). Sounds were calibrated in intensity at the location of the mouse ear using a probe microphone (Bruel & Kjaer).

For V1 recordings, auditory only stimulations were performed in two different contexts: either in complete darkness (screen turned off in the sound and light isolated box enclosing the microscope) or in dim light (screen turned on with black background, luminance measured to 0.57 candela per square meter).

### Intrinsic optical imaging recordings

To localize the calcium-imaging recordings with respect to the global functional organization of the cortex, we performed intrinsic optical imaging experiments under isoflurane anaesthesia (1%). The brain and blood vessels were illuminated through the cranial window by a red (intrinsic signal: wavelength 780 nm) or a green (blood vessel pattern: wavelength 525 nm) light-emitting diode.

To localise the visual cortex, the reflected light was collected at 15 Hz by a charge-coupled device (CCD) camera (Foculus IEEE 1394) coupled to the epifluorescence light path of the Femtonics microscope (no emission or excitation filter). A slow drifting bar protocol was used: a white vertical bar was drifting horizontally over the screen width for 10 cycles at 0.1Hz, from left to right in half of the trials, and from right to left in the other half. After band-passing the measured signals around 0.1Hz, and determining their phase in each pixel and for each condition, both the preferred bar location and the hemodynamic delay could be determined for each pixel, yielding azimuth maps as in (**Fig 2e**). Similarly, elevation maps were obtained in a subset of the animals using a horizontal bar drifting vertically. These maps coincided with those obtained in previous studies^38^, so we could determine V1 border as the limit between pixels displaying and not displaying retinotopy tuning.

To localise the auditory cortex, the reflected light was collected at 20 Hz by a CCD camera (Smartek Vision, GC651M) attached to a custom-made macroscope. The focal plane was placed 400 mm below superficial blood vessels. A custom-made Matlab program controlled image acquisition and sound delivery. We acquired a baseline and a response image (164 × 123 pixels, ~3.7 × 2.8 mm) corresponding to the average images recorded 3 s before and 3 s after sound onset, respectively. The change in light reflectance (ΔR/R) was computed then averaged over the 20 trials for each sound frequency (4, 8, 16, 32 kHz, whitenoise). A 2D Gaussian filter (σ = 45.6 µm) was used to build the response map and determine the localisation of the auditory cortex due to the stereotypical response map produced by the 4kHz sound. Sounds were trains of 20 white noise bursts or pure tone pips separated by 20 ms smooth gaps.

### Eye tracking and trial filtering

Left eye measurements (eye size, pupil diameter, pupil movement) were measured by tracking the eye using a CCD camera (Smartek Vision, GC651M). A Python software was used to capture images from the camera at 50Hz, synchronized with the cortical recordings.

These movies were analyzed off-line using custom automatic Matlab programs that traced the contours, first of the eyelid, second of the pupil. The eye lid shape was approximated by two arcs and involved the estimation of 6 parameters (4 for the coordinates of the two points where the arcs join, 2 for the y-coordinates of the crossings of the arcs with the vertical line at halfway between these two points; see **Fig 2c**, bounds for the parameters were set by hands and appear in yellow). The pupil shape was approximated by an ellipse, described by 4 parameters (center x and y, radius and eccentricity). Both estimations were performed by maximizing the difference between average luminance inside and outside the shape, as well as the luminance gradient normal to the shape boundary; in addition they were inspected manually and corrected occasionally inside a dedicated graphic interface. In the "dim light" and "bimodal" (but not the “dark”) contexts, it was observed that saccades correlated with population activity increase (e.g. **Supplementary Fig. 2**), therefore we discarded all trials from these contexts displaying saccades. To do so we computed the maximal distance between pupil center positions at different instants during the 2 seconds of stimulation 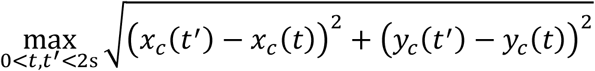 and discarded trials where the change in gaze exceeded the mouse visual acuity of 2°, i.e. where this distance exceeded 57.6µm (assuming an eye radius of 1.65mm).

### Calcium imaging data summary

The data obtained from the different imaging experiments consisted of the following.

A1 recordings: 3,771 cortical neurons from 18 L2/3 imaging areas in 7 animals, bimodal context only. And 1,593 neurons projecting to V1 from 14 L2/3 areas in 3 additional animals.

V1 recordings: 18,925 neurons from 35 L2/3 imaging areas in 9 mice, and included the three “dark”, “dim light” and “bimodal” blocks in random order. Eye tracking was performed in 23 of these sessions (9,849 neurons, 7 mice; only these sessions where used for the analyses of the “dim light” and “bimodal” context, which required filtering out trials with saccades).

V1 recordings with labelling of Gad-positive interneurons: an additional 4,348 neurons from 12 L2/3 imaging sessions (depth ranging from 140 to 300 microns), and 265 GAD2 positive neurons from 7 L1 imaging sessions (depth from 30 to 100 microns). All of these sessions had eye tracking, and included the “dark” and “dim light” blocks. 6 L2/3 sessions also included “bimodal” blocks, whereas the 13 remaining sessions included a second “dim light” block where the left (contralateral) eye was covered by a dark mask (**Fig. 3f**).

### Calcium imaging pre-processing

Data analysis was performed with custom-made Matlab scripts available upon request. Every frame recorded was corrected for horizontal motion to a template image using rigid body registration (all sessions with visible z motion were discarded). Regions of interest were then automatically selected and human checked as the cell bodies of neurons with visually identifiable activity^35^ and the mean fluorescence signal F(t) was extracted for each region. We also estimated the local neuropil signal Fnp(t) for each neuron^27^ and subtracted a fraction of it from the neuron signal. This fraction was set to 0.7, according to a previous calibration for GCAMP6s in mouse visual cortex^34^. Baseline fluorescence F_0_ was calculated as the minimum of a Gaussian-filtered trace over the 5 neighbouring 5s imaging blocks and fluorescence variations were computed as f(t) = (Fc(t) - Fc0)/Fc0n. Analyses were performed either on these normalized fluorescence signals, or on estimations of the firing rate obtained by temporal deconvolution^36,37^ as r(t) = f’(t) + f(t)/τ, in which f’(t) is the first derivative of f(t) and τ = 2 s, as estimated from the decays of the GCAMP6s fluorescent transients ^34^. This simple method efficiently corrects the strong discrepancy between fluorescence and firing time courses due to the slow decay of spike-triggered calcium rises^37^, and was preferred to more advanced deconvolution methods because it does not bias deconvolution towards absence of activity (i.e. interpreting small signal as noise) in cells displaying poor signal to noise ratio. However it does not correct for the relatively slow rise time of GCAMP6s, producing a time delay on the order of 70 ms between peak firing rate and peak deconvolved signal^27^.

### Data analysis

Data analysis was performed with custom-made Matlab and Python scripts available upon request.

Clustering of auditory responses in the auditory and visual cortices (**Fig. 1**), and of bimodal conditions in the visual cortex (**Fig. 6**) was performed using the following procedure. Calcium responses were deconvolved, baseline subtracted, blank subtracted, averaged over trials, and the responses to all considered stimuli were concatenated. Hierarchical clustering was performed using the Euclidean metric and Ward method for computing distance between clusters. To determine the number of clusters we moved down the clustering threshold of until clusters became redundant (overclustering) as assessed visually. This method clusters neurons irrespective of whether they significantly responded to the stimuli. As a large number of neurons were not (or very weakly responsive) to the stimuli, a large fraction of the neurons were grouped into “noisy” clusters displaying no obvious sensory response. These “noisy” clusters were eliminated based on their noise levels measured as the mean distance between the raw average responses in the cluster and their time-filtered version (Gaussian kernel, σ = 95ms). All clusters with less than 5 neurons or with a noise level larger than 20% (ΔF/F.s^−1^) were discarded. These thresholds were chosen by visual inspection of the obtained clusters. To make sure cell type distributions were not skewed by the fact that clustering outputs the most robust auditory responses (Fig. 1), we re-aggregated neurons discarded as nonresponsive if the mean correlation of their activity signature with any of the identified clusters was larger than 0.2. This procedure did not change the mean cluster response profiles and did not qualitatively impact the conclusions drawn on the distributions of the different clusters in V1-projecting and L2/3 neurons. In fact, the fraction of cells preferring down-ramps in V1-projecting non-re-aggregated clusters was even larger (71% in V1-project for 37% in L2/3, 4-fold difference for loud onset neurons), indicating that the observed preferences are even clearer in strongly driven neurons.

### Statistics

Data are displayed as mean and S.E.M.

To assess the significance of the relative distribution of functional cell types in V1-projecting neurons and in layer 2/3 neurons, we performed a bootstrap analysis. The null-hypothesis is that both populations have the same distribution of clusters. To simulate this hypothesis we pooled together all neurons and performed 10,000 random partition of the pool population into the number of V1-projecting and L2/3 neurons. For each partition, we computed the difference of between the fractions of each clusters across the two partitions. This led to distributions of expected fraction differences for each cluster under the null hypothesis. The p-value was computed from the percentile of which the actual observed fraction difference was located.

Significant responses for individual neurons were detected using the non-parametric Wilcoxon rank-sum test. Raw calcium fluorescence responses were subtracted for prestimulus level, and averaged over a time window near the response peak. The vector of such responses for different trials was compared to the same computations performed on blank trials (unless responses to two different conditions were compared, such as auditory responses in the dark vs. in a dim light). Unless specified otherwise, we used a statistical threshold of 1% for detecting responding neurons; by definition this means that on average 1% of the neurons *not responding* to the condition would be detected as responding (false positive). Therefore in all the histogram displays of the fractions of responding neurons (e.g. **Fig. 5c, f**) we masked the first 1% as being potentially only false positive detections.

To assess the significance of supra- or sub-linear responses to audio-visual combinations in individual clusters resulting from the clustering of bimodal responses, we used a bootstrap consisting in shuffling the different trial repetitions. The null hypothesis is that the cluster’s average bimodal responses can be predicted as the sum of average visual and auditory responses, after baseline has been subtracted. To test this hypothesis, we first computed, for each neuron, individual “repetitions” of the linear predictions by adding responses to one visual presentation and to one auditory presentation. Then we performed 1e6 shufflings of the labels of “linear prediction” and “bimodal response” trials, yielding a distribution of the expected difference between their averages under the null hypothesis. The p-values were computed from the percentile in which the actual nonlinear difference was located.

### Model

Simulations were performed using a rate model with 2 or 3 populations and no synaptic delay. The spiking activity *r_i_* of population *i* followed the equation:

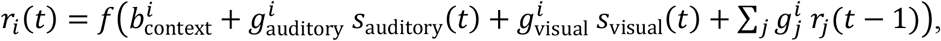

Where *s*_auditory_ and *s*_visual_ are the auditory and visual input (displaying an exponentially decaying signal and a plateauing signal, respectively), 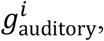 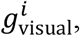 and 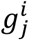 the connectivities to input and to the other populations (see model schematics in **Fig. 4a and Supplementary Fig. 4** for which connections are non-zero). 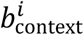 is a baseline accounting for other inputs, and whose value can depend on the context (“dark” or “light”). *f* is a nonlinear function consisting of two linear segments (**Fig. 4a**) that accounts for spiking threshold or further nonlinear (e.g. dendritic) computations. The first segment is not a constant zero accounts for the fact that the other inputs summarized in 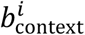 are in fact stochastic, and can lead the cell to fire even when its average potential is below threshold.

### Behavior

We tested the effect of playing sounds during a visual detection task. Water-deprived mice (33 ml g^−1^ per day) were head-fixed and held in a plastic tube on aluminium foil. The first day they were habituated to the fixation for one hour and manually rewarded. Then they were trained to lick when the rewarded stimulus (S+), the 2 second either looming or receding white disk on dark background, was presented. The mouse had to produce at least one lick on a stainless steel water spout to receive a 5 ml water drop. Licks were detected by changes in resistance between the aluminium foil and the water spout. To refrain mice from constantly licking, trials were started only following a period of 3 seconds without any lick. During the first sessions the reward was delivered immediately following the first lick; later the reward was delivered at least 1.7s after stimulation onset. After mice successfully responded to S+, achieving a steady state level of performance (6 sessions of ~200 episodes), up- or down-ramping sounds were introduced and were played together with visual stimulation for half of the episodes; reward remained the same on these bimodal conditions.

## Data availability

The data that support the findings of this study and the analysis code are available from the corresponding author upon request.

## Author contributions

TD, AK and BB designed the study, performed and analyzed the experiments, and wrote the paper.

## Acknowledgments

We thank A. Fleischmann, Y. Frégnac and J. Letzkus for helpful discussions and comments on the manuscript. We thank A. Fleischmann for providing CAV2-Cre viruses. This work was supported by the Agence Nationale pour la Recherche (ANR “SENSEMAKER”), the Marie Curie FP7 Program (CIG 334581), the Human Brain Project (SP3 - WP5.2), the Fyssen foundation, the DIM “Region Ile de France”, the International Human Frontier Science Program Organization (CDA-0064-2015), the Fondation pour l’Audition (Laboratory grant) and the Paris-Saclay University (Lidex NeuroSaclay, IRS iCode and IRS Brainscopes).

